# Diversification of CD1 molecules shapes lipid antigen selectivity

**DOI:** 10.1101/2020.11.10.377556

**Authors:** Nicole M. Paterson, Hussein Al-Zubieri, Matthew F. Barber

**Affiliations:** Institute of Ecology and Evolution, University of Oregon, Eugene, OR 97403; Department of Chemistry and Biochemistry, University of Oregon, Eugene, OR 97403; Department of Biology, University of Oregon, Eugene, OR 97403

## Abstract

Molecular studies of host-pathogen evolution have largely focused on the consequences of variation at protein-protein interaction surfaces. The potential for other microbe-associated macromolecules to promote arms race dynamics with host factors remains unclear. The cluster of differentiation 1 (CD1) family of vertebrate cell surface receptors plays a crucial role in adaptive immunity through binding and presentation of lipid antigens to T-cells. Although CD1 proteins present a variety of endogenous and microbial lipids to various T-cell types, they are less diverse within vertebrate populations than the related major histocompatibility complex (MHC) molecules. We discovered that CD1 genes exhibit a high level of divergence between simian primate species, altering predicted lipid binding properties and T-cell receptor (TCR) interactions. These findings suggest that lipid-protein conflicts have shaped CD1 genetic variation during primate evolution. Consistent with this hypothesis, multiple primate CD1 family proteins exhibit signatures of repeated positive selection at surfaces impacting antigen presentation, binding pocket morphology, and TCR accessibility. Using a molecular modeling approach, we observe that inter-species variation as well as single mutations at rapidly-evolving sites in CD1a drastically alter predicted lipid binding and structural features of the T-cell recognition surface. We further show that alterations in both endogenous and microbial lipid binding affinities influence the ability of CD1a to undergo antigen swapping required for T-cell activation. Together these findings establish lipid-protein interactions as a critical force of host-pathogen conflict and inform potential strategies for lipid-based vaccine development.

## Introduction

Early detection of pathogen-specific molecules by the immune system can mean the difference between resistance, latency, or succumbing to infectious disease. Previous studies have illustrated that host-pathogen protein interaction surfaces are hotspots for repeated natural selection by conferring resistance or susceptibility to infection (van der Lee et al. 2017; Daugherty and Malik 2012; Enard et al. 2016). Such conflicts between hosts and pathogens can give rise to a variety of evolutionary dynamics including Red Queen arms races (Daugherty and Malik 2012; Van Valen 1973), frequency-dependent selection (Takahata and Nei 1990), and over-dominance (Hughes and Nei 1990; Takahata and Nei 1990; Nei and Rooney 2005). While vertebrate immune systems are tuned to recognize a wide variety of pathogen-associated macromolecules including DNA, RNA, lipids, and glycans, our understanding of host-pathogen evolutionary conflicts is largely restricted to protein-protein interactions (Sawyer et al. 2004; Barber and Elde 2014; Choby et al. 2018; Elde et al. 2009). In the case of lipid and lipopeptide antigens, the production of a functional molecule involves the synthesis of precursors that are further processed by enzymatic modifications. As such, evolutionary dynamics involving these macromolecules and their host receptors may be distinct from cases of protein-protein co-evolution.

The major histocompatibility complex (MHC) superfamily comprises a variety of cell surface proteins which present self and foreign antigens to T-cells. Recognition of foreign antigens by T-cell receptors (TCRs) leads to T-cell activation and initiation of an adaptive immune response (Frank 2002). Multiple evolutionary forces are hypothesized to contribute to the immense diversity in MHC haplotypes, including over-dominance wherein increased heterozygosity is favored by selection and polymorphisms are maintained over time (Takahata and Nei 1990). In addition to class I and class II MHC molecules which present peptide antigens, the related cluster of differentiation 1 (CD1) and MR1 molecules have been shown to present lipid and lipoprotein antigens to T-cells (Barral and Brenner 2008; Birkinshaw et al. 2015; Blumberg et al. 1995; Mori, Lepore, and De Libero 2016; Zajonc and Flajnik 2016). CD1 molecules display rare and infrequent polymorphism (Golmoghaddam et al. 2013; Han et al. 1999), with limited genetic diversity within humans and other populations relative to class I and II MHC. The MHC and CD1 gene families therefore appear to have experienced divergent evolutionary paths after their duplication from a common ancestor, with respect to antigen recognition and population genetic variation. CD1 paralogs are divided into groups, whereby group 1 CD1 family members (human CD1a, CD1b, and CD1c) present antigen primarily to cytotoxic CD8+ T-cells (Mori, Lepore, and De Libero 2016, 1). Group 2 CD1 molecules (CD1d in humans) present antigen to invariant natural killer (iNK) T-cells (Pereira and Macedo 2016).

CD1 and MHC arose in jawed fishes with the advent of the B-cell and T-cell immune receptors during the genesis of the adaptive immune system of vertebrates (Barral and Brenner 2008). After the gene duplications that gave rise to ancestral MHC and CD1, vertebrate CD1 paralogs expanded in through repeated duplication events (Dascher 2007). This initial expansion was followed by differential pseudogenization and expansion of CD1 paralogs to various degrees across vertebrate species (Rogers and Kaufman 2016). As such, CD1 gene content varies widely across vertebrates: primates have a single copy of the CD1a paralog, mice possess none, dogs encode six, and horses possess five (Dascher 2007). In primates, CD1d is believed to represent the most deeply conserved member of the CD1 family (Salomonsen 2007.). CD1d receptors can display antigen to specialized iNKT-cells, which are able to mount an earlier response to infection due to their dual role in innate as well as adaptive immunity (Pereira and Macedo 2016). Current evidence indicates that human CD1e does not present antigen (Garcia-Alles et al. 2011) but rather assists in antigen loading onto CD1d in lysosomes and endosomes (Cala-De Paepe et al. 2012). This wide range of responsive T-cell types, along with evidence that these non-classical T-cell types mount an early immune response to infection (Godfrey et al. 2015), makes human CD1-expressing cells surprisingly flexible responders to infections even with their lack of exceptional sequence variation.

CD1 molecules possess an extracellular domain containing a sub-surface hydrophobic binding pocket used to present antigen to CD1-restricted T-cells (Figure 1B) (Barral and Brenner 2007). During the adaptive immune response, CD1 on the surface of antigen-presenting cells activate T-cells by displaying specific classes of hydrophobic ligands to T-cell receptors (Figure 1C) (Blumberg et al. 1995; Barral and Brenner 2008; Chancellor, Gadola, and Mansour 2018). According to structural studies, CD1a has the smallest of the human CD1 binding pockets with a volume of about 1280 angstroms (Ly and Moody 2014). After the gene duplication event that gave rise to this paralog, CD1a likely evolved to present either self-lipids or small exogenous lipopeptides (Mori, Lepore, and De Libero 2016). Consistent with this hypothesis, CD1a has been crystallized in complex with self-lipids sphingomyelin, lysophosphatidylcholine (Birkinshaw et al. 2015), sulfatide (Zajonc et al. 2003), as well as the mycobacterial lipopeptide analog didehydroxymycobactin (Zajonc et al. 2005). The binding pocket of human CD1a is composed of a double chambered cavity termed the A’ and F’ pockets with a single (A’) portal that coordinates the presentation of lipid antigen to the TCR (Figure 1B) (Zajonc et al. 2005). The TCR lands just above the A’ pocket on a surface termed the A’ roof (Zajonc et al. 2005). The diminutive size of the CD1a A’ pocket is thought to be formed by the electrostatic interaction of the two side chains belonging to the A’ roof that also draw the two parallel alpha helices of the pocket in close proximity, while an amino acid sidechain blocks the base of the pocket thereby limiting size of tail groups that can be accommodated (in human CD1a this amino acid is valine 28) (Zajonc et al. 2003). Several other CD1 homologs (except for CD1c) lack this roof structure (Blumberg et al. 1995). CD1a does not feature a late endosomal targeting element and does not require low pH for antigen binding as is the case for other CD1 proteins such as CD1b, CD1c and CD1d (Chancellor, Gadola, and Mansour 2018).

**Figure 1.**
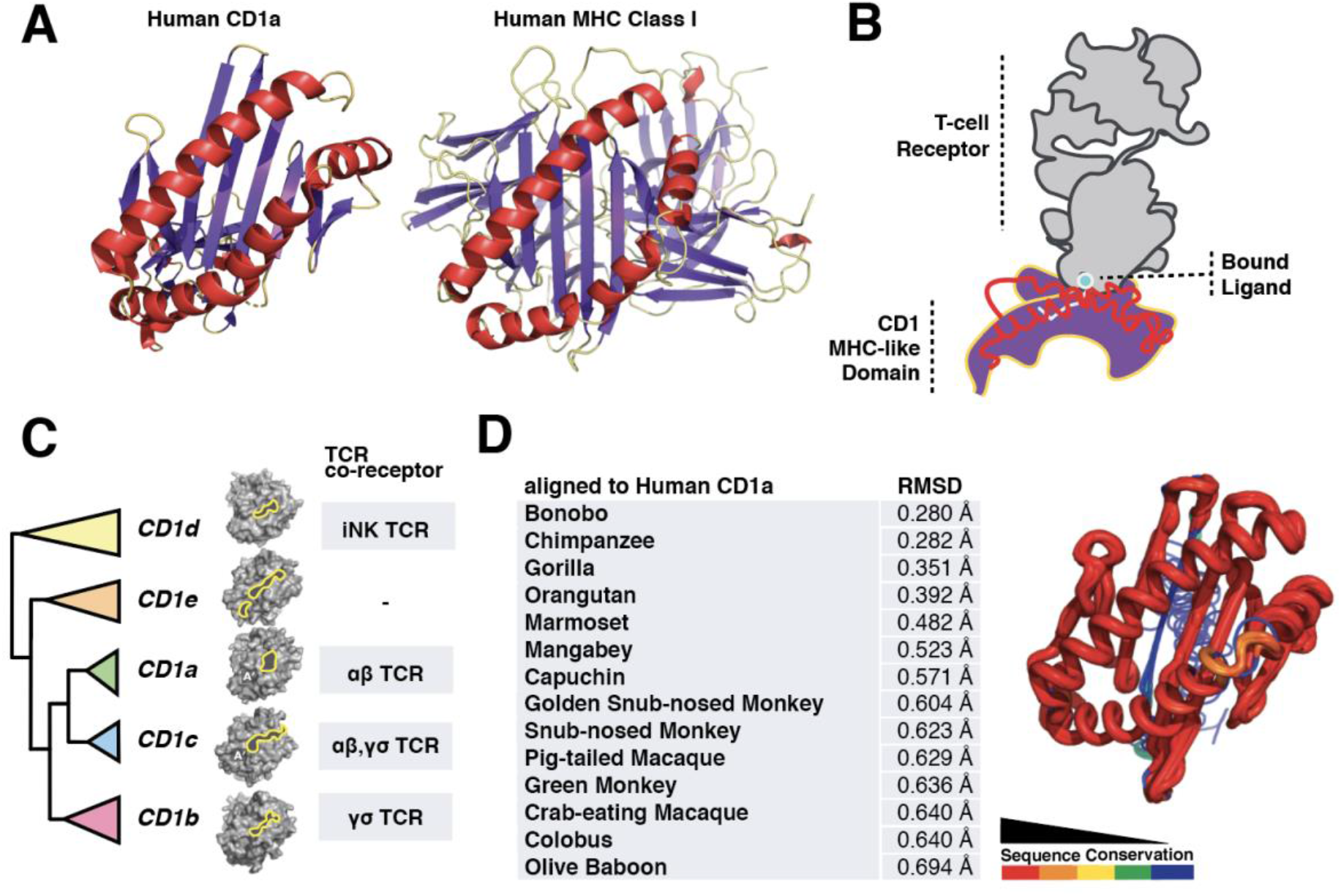
Diversity of the CD1 gene family in primates. **(A)** Ribbon diagrams of human CD1a (PDB ID: 5J1A) and MHC (PDB ID:1AKJ) with alpha helices highlighted in red, beta-sheet in purple, loops in yellow. **(B)** Illustration of the CD1-TCR interaction where CD1 bind lipid tails in a hydrophobic pocket with polar head groups typically exposed. The TCR (grey) “reads” the displayed antigen leading to T-cell activation. **(C)** Cladogram representing phylogenetic relationship of primate CD1a-e paralogs used in this study with surfaces generated in PyMol and antigen binding pockets outlined in yellow (PDB IDs: 5J1A, 4ONO, 4MQ7, 3S6C). TCR types that recognize each CD1 paralog are also indicated. **(D)** Primate CD1a diverges most in the lipid binding domain, which may alter pocket morphology and TCR interactions. Most of the sequence divergence in the primate CD1a proteins is predicted to exist in the beta-sheet that transects the center of the protein with some variation in the central surface region.

The immense diversity of the MHC family within and between populations at surfaces necessary for peptide antigen recognition have made these genes classic study systems of adaptive protein evolution (Grimholt 2016; Danchin and Pontarotti 2004; Castro, Luoma, and Adams 2015). CD1 molecules possess similar structure and function to class I and II MHC proteins, although their relative lack of diversity at the population level has been attributed to a lack of diversity in their cognate lipid ligands. While variation in pathogen-derived lipids has been implicated in host immune recognition and virulence (Chandler et al. 2020), the potential for lipid antigens to promote evolutionary arms races with hosts is unclear. In the present study we used the CD1 family as a system to investigate diversity and evolution of lipid antigen recognition by the vertebrate immune system.

## Results

### Diversification of the CD1 gene family in primates

A comparison of MHC class I and CD1 protein structures illustrates the homology between these antigen presentation molecules (Figure 1A). CD1 presents antigen to the TCR with the lipid tail groups tucked into the hydrophobic pocket and head groups exposed where they are “read” by the TCR (Figure 1B). Distinct CD1 molecules present antigen to a wide variety of T-cell types (Figure 1C) (Godfrey et al. 2015). To assess patterns of genetic diversity among primate CD1 family members, we first assembled a collection of CD1 paralogs across simian primates from publicly available genome databases and generated a phylogenetic gene tree using PhyML (Figure 1C, Supplemental Figure S1). The five human CD1 paralogs are largely conserved across primate species, allowing us to reliably assess structural and genetic diversity within this family.

A comparison of the sequences among CD1 paralogs revealed a striking degree of diversification, particularly in the MHC-like domain responsible for lipid antigen presentation (Figure 1D). To assess the potential consequences of this variation on CD1 function, we plotted the structural conservation among primate CD1a orthologs using color by conservation (Mura et al. 2010) (Figure 1D). Our analysis revealed several hotspots of high amino acid divergence among CD1 molecules, focused on both interior regions of the antigen binding pocket as well as surface helices that are known to contact the TCR. Together our results indicate that, despite their limited polymorphism within populations, CD1 paralogs exhibit a surprising degree of genetic divergence between simian primate species.

### Signatures of repeated positive selection acting on primate CD1 genes

Given their relative lack of within-population diversity, we were surprised by the elevated genetic divergence between primate CD1 orthologs. We hypothesized that this variation could be the result of repeated positive selection in response to variation in lipid antigen structures. To identify potential amino acid sites that may have been subject to repeated positive selection, we used the codeml package from PAML (Yang 2007) in addition to MEME (Murrell et al. 2012) and FuBar (Murrell et al. 2013) from the HyPhy software package to detect elevated dN/dS (ω) at sites within CD1 gene paralogs across primates. Elevated ω values were detected for all CD1 family members (Figure 2A) with the exception of CD1b, consistent with elevated nonsynonymous substitution rates associated with positive selection. We noted that the majority of rapidly-evolving sites among CD1 paralogs were focused in the MHC-like domain which is responsible for lipid binding (Figure 2A). These results suggest that multiple members of the CD1 family have undergone repeated episodes of positive selection in simian primates specifically within regions important for lipid antigen presentation.

**Figure 2.**
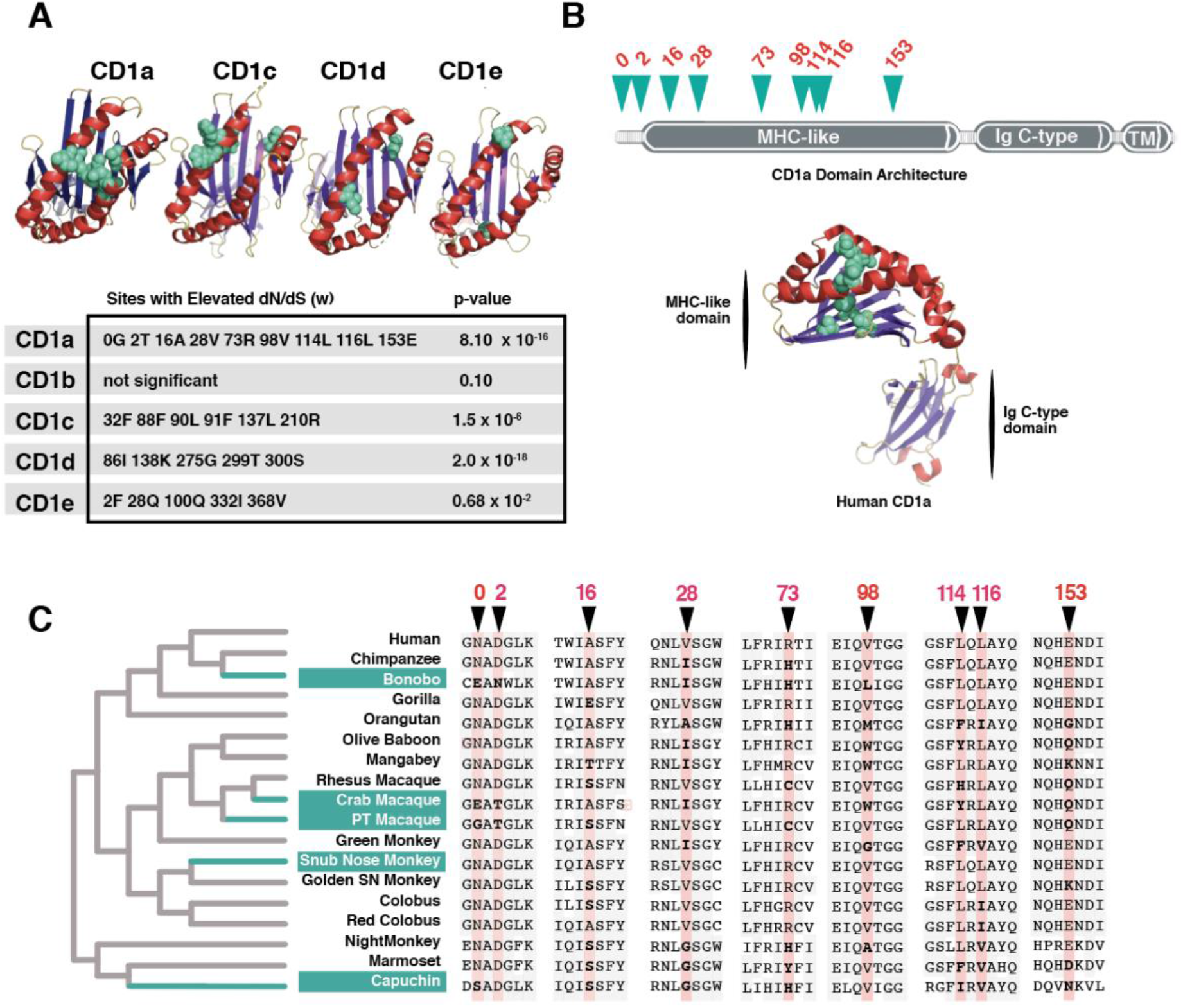
Evidence of repeated positive selection among primate CD1 orthologs. **(A)** Amino acid sites exhibiting strong signatures of positive selection (elevated dN/dS) are highlighted in teal and mapped onto corresponding crystal structures. Alpha helices are denoted in red, beta-sheet in purple. (PDB IDs: 5J1A, 4ONO, 4MQ7, 3S6C). Table summarizes positions in CD1 paralogs contributing to signatures of positive selection as well as statistics from PAML M7-M8 model comparisons. **(B)** Sites with elevated dN/dS indicative of positive selection (teal) cluster in the MHC domain of CD1a protein (PDB ID: 5J1A). Alpha helices denoted in red, beta-sheet in purple. **(C)** Multiple sequence alignment of primate species used to calculate dN/dS ratios for CD1a paired with phylogenetic species tree highlighting the branches (teal) predicted by aBSREL to be undergoing episodic positive selection.

Having detected evidence of positive selection acting on multiple CD1 family members, we chose CD1a for additional in-depth analysis. CD1a has less stringent lipid loading requirements than other CD1 homologs as it does not require a reduced pH environment encountered in late endosomes nor does it have a known adapter protein required for antigen loading (Barral and Brenner 2007). For these reasons, we anticipated that empirical and molecular modeling studies of antigen recognition complex be less complex for CD1a than other paralogs. CD1a has been shown to present mycobacterial antigens from the cell surface where it interacts with langerin on Langerhans cells (Mizumoto and Takashima 2004), a specialized dendritic cell type that surveys epithelial monolayers for molecular indicators of infection.

Seeking to determine what domains of CD1a are subject to positive selection, we mapped the sites with high ω values from our previous analysis. Results show clustering of rapidly evolving sites in the MHC-like domain, similar to those observed with other CD1 paralogs (Figure 2B, Supplemental Figure S4). These sites also map to regions where CD1a is likely to interface with lipid antigen or the TCR, suggesting that selection may have acted to alter lipid binding and T-cell interactions. The majority of variable sites between CD1a orthologs cluster to a region of the protein near the center of the binding pocket and around the outer surface (Figure 2B). We grouped all of the rapidly-evolving sites we identified in this study into three categories: residues located at or near the TCR landing site (the A’ roof in the human structure), residues within the binding pocket, and residues in the N-terminus for which we have no structural information. Overall, predicted structural features do not correlate well with phylogenetic relatedness, consistent with multiple lineages undergoing episodic selection (Figure 2C, Figure 3A).

**Figure 3.**
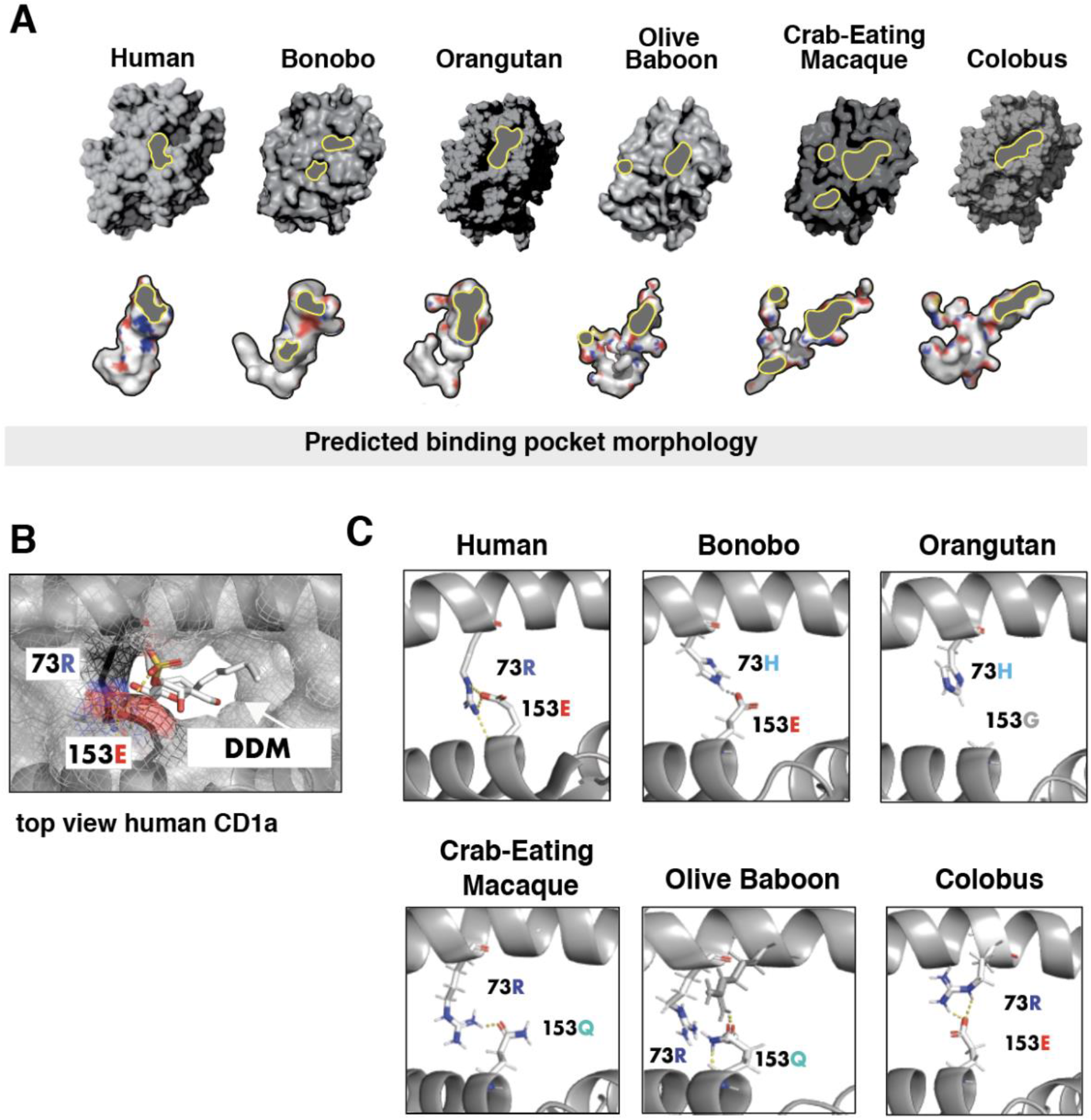
Structural modeling illustrates diversity at the CD1a T-cell interaction interface. **(A)** Predicted surface attributes of various primate CD1a structures. Surface characteristics across selected primates reveals differences in portal size, number of portals and pocket morphology. Portals where T-cell receptor “reads” head group are highlighted with grey/yellow outlines. Pocket morphologies and electrostatic properties are shown below surface models. **(B)** PyMol generated top-view of human CD1a bound to dideoxy-mycobactin (PDB ID:1XZO) to show importance of rapidly-evolving positions 73 and 153 in coordinating head groups of antigenic ligand. Note hydrogen bonding between head group and 153E. **(C)** New and Old World primate CD1a at A’ Roof where CD1a interacts with TCR. Notably, the orangutan model does not form roof structure due to mutation at site under selection. Olive baboon and crab-eating macaque form A’ roof with residue of differing property at site 153.

### Accelerated evolution of the CD1a-TCR interface

To assess how variation in CD1a may influence immune functions in primates, we used I-TASSER to generate predicted structures of several CD1a orthologs (Figure 3A). Of the hominoid structures modeled, human, bonobo, and orangutan have remarkably different topologies at the TCR-CD1a interface as well as the geometry of the internal binding pocket (Figure 3A). The morphologies of the binding pockets vary widely, most notably in the crab-eating macaque which is predicted to contain one main and two accessory portals, with a narrow meandering channel (bottom Figure 3A). The length and volume of the pockets limits the types of lipid tail groups that can be accommodated, while the size and location of the portals has effects on how well the T-cell receptor can read the antigen presented (Birkinshaw et al. 2015).

In human CD1a, the A’ roof is hypothesized to aid in determining whether an antigen will elicit an immune response by supporting interactions with the TCR and assisting in display of the ligand head-group (Figure 3B) (Birkinshaw et al. 2015; Zajonc et al. 2005). The predicted orangutan CD1a structure lacks an A’ roof entirely (Figure 3C), while bonobo possesses two portals. Additionally, it has been speculated that disruption of hydrogen bonding between R73, R76, and E154 that form the A’ roof may indicate whether a given ligand will stimulate TCR activation (Birkinshaw et al. 2015). However, several of the CD1a structures are predicted to form an A’ roof that does not depend on this particular interaction. For example, crab-eating macaque and olive baboon CD1a are predicted to form a relatively unique A’ roof composed of an R73/153Q linkage that does not involve R76 (Figure 3C). The TCR does not recognize CD1a-bound ligands without adequate projection of lipid head groups, and it is likely that hydrogen bonding between the head groups of smaller ligands and residues that make up the portal are important for display. Headless ligands buried in the CD1a pocket, for example, can result in T-cell auto-reactivity (de Jong et al. 2010). Site 153, which is remarkably variable across primates (Figure 2D) has been shown to form a hydrogen bond in human CD1a to the head group of self antigen lysophosphatidylcholine and sulfatide in addition to its role in forming the A’ roof (Zajonc et al. 2003; Birkinshaw et al. 2015). This site bears a glycine in orangutan, with no ability to form an A’ roof or salt bridges with ligand (Figure 3C). Altogether, these predicted structural differences suggest that natural selection may have had a significant impact on the ability of CD1a to display self or foreign lipid antigens across related primates.

### Structural remodeling of the CD1a antigen binding pocket

To determine whether the structural differences observed across CD1a paralogs are likely to have functional consequences for antigen recognition, we applied a ligand-docking approach using AutoDock Vina. We were particularly interested to test affinity differences between endogenous and exogenous lipid ligands, since the current model for CD1a antigen presentation is a swapping mechanism wherein lower affinity endogenous ligand is replaced at the cell surface by higher affinity exogenous ligand. We used ligands previously crystallized in complex with human CD1a in our studies since there is a wealth of structural information available on these particular binding interactions. In our docking simulations, we found that a single loop region is required to re-dock all ligands in the CD1a binding pocket. We assigned flexibility to this region for all the structures tested, as well as any non-bonded side chains in the region of the portal (Supplemental Figure S2). We believe this is likely the region responsible for conferring flexibility in the native CD1a, which must be flexible enough to accommodate ligands with a diversity of molecular weights (molecular weight of urushiol is 330 g/mol, dideoxy-mycobactin is 838 g/mol).

We next measured predicted CD1a binding affinities for the panel of lipid ligands including endogenous ligands sphingomyelin, sulfatide, lysophosphatidylcholine and exogenous ligand dideoxy-mycobactin (Figure 4A, Supplemental Figure S6). Given that CD1a is believed to swap endogenous lipids for exogenous lipids based on differences in relative binding affinities, we then estimated the likelihood of a lipid-swap using our panel of ligands. We calculated the fold differences in Kd between the highest affinity endogenous lipid and compared this to the Kd of the exogenous ligands. Crab-eating macaque, snub-nosed monkey, olive baboon, capuchin, mangabey, bonobo, and human were predicted to swap out endogenous for mycobacterial exogenous ligand dideoxy-mycobactin (DDM) (Figure 4B). It is worth noting that we make the assumption that the endogenous lipid with the lowest Kd is also present in abundance, which we cannot not know for certain *in vivo*. It has been shown in previous studies that human CD1a molecules bind to a diverse repertoire of lipid types *in vitro* (Birkinshaw et al. 2015). Since lipid profiles aren’t available for all cell types in the primates we studied, we chose this simplification as a rough estimate for feasibility of lipid swapping.

**Figure 4.**
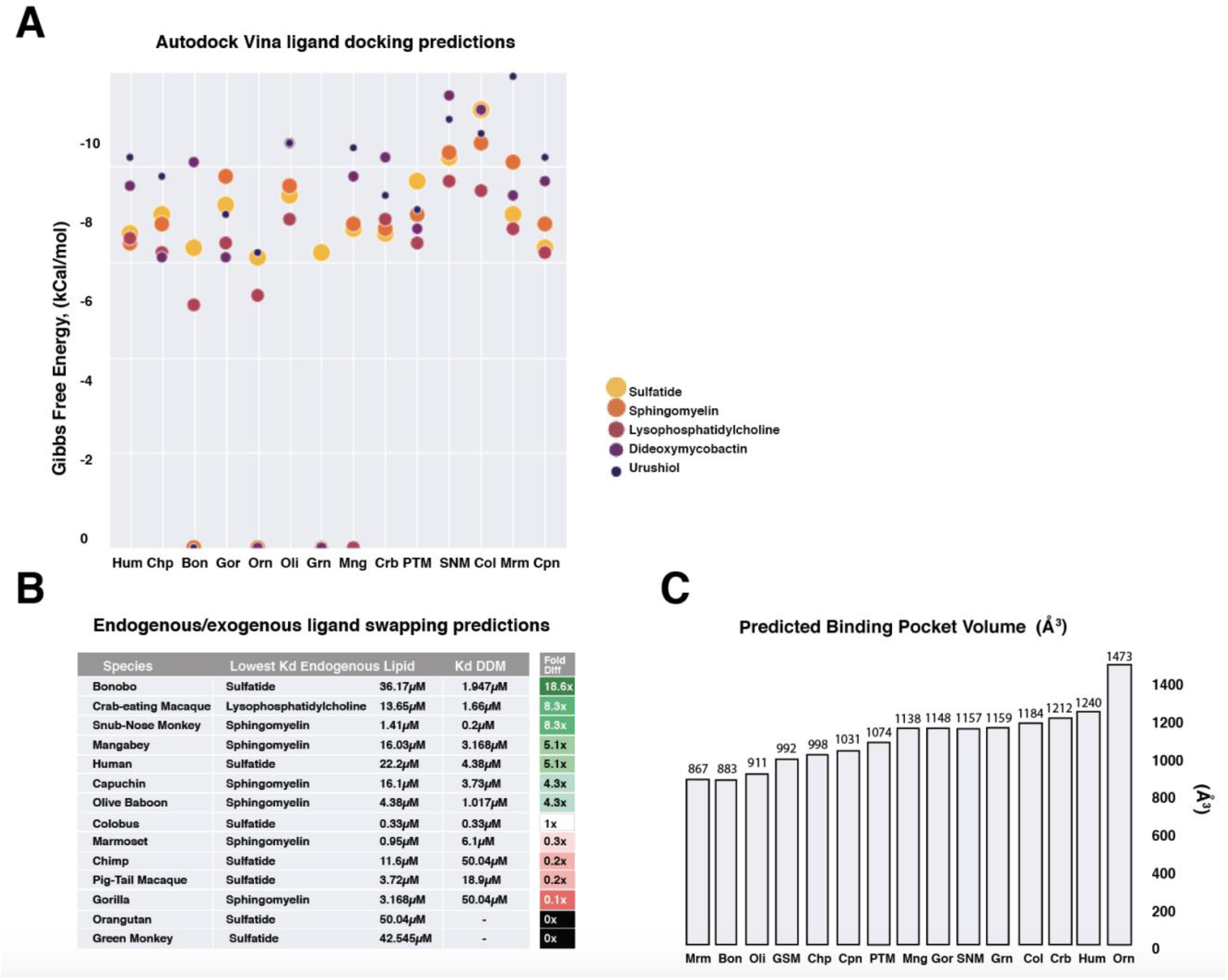
Divergence of CD1a shapes predicted endogenous and exogenous lipid antigen affinities. **(A)** Table plotting relative Gibbs free energy values for all ligands tested by ligand docking predictions using AutoDock Vina. Lowest energy values for each set are plotted. Sulfatide, sphingomyelin, lysophosphatidylcholine are endogenous lipid ligands. Dideoxymycobactin is a synthetic lipid analog of *M. tuberculosis* siderophore mycobactin. Urushiol is the etiological agent of poison ivy rash. **(B)** Lipid-swapping predictions based on predicted Kd from docking studies. Great apes highlighted in blue, Old World monkeys are highlighted in grey and New World monkeys in turquoise. **(C)** Predicted pocket volume differences vary most in great apes and do not correlate with phylogenetic relationships. Legend: **Hum**-Human, **Chp**-Chimpanzee, **Bon**-Bonobo, **Gor**-Gorilla, **Orn**-Orangutan, **Oli**-Olive Baboon, **Grn**-Green Monkey, **Mng**-Mangabey, **Crb**-Crab-eating macaque, **SNM**-Snub-nosed monkey, **Col**-Colobus, **Mrm**-Marmoset, **Cpn**-Capuchin

A notable result from the ligand docking predictions shows that binding profiles failed to group by species phylogeny consistent with branch-site test results that detected several branches undergoing multiple bouts of episodic positive selection (Supplementary Figure S5). Unlike in humans where the largest binding pocket also has the most promiscuous ligand binding profile, ligand docking predictions do not group higher affinity binding with predicted pocket volume (Figure 4C). These findings indicate that predicted structural alterations in the CD1a ligand binding pocket have significant impacts on recognition of both endogenous and pathogen-derived antigens which collectively shape downstream T-cell activation.

### Modeling the effects of rapidly-evolving sites on antigen presentation by CD1a

We next assessed how variation at single rapidly-evolving positions in the CD1a binding pocket may alter lipid antigen recognition. Using PyMol, we substituted single extant amino acids for the ancestral amino acid at sites undergoing positive selection (ancestral sites predicted by DataMonkey package SLAC (Pond and Frost 2005)) and used these altered structures in our ligand-docking simulation. We first tested the effects of selected sites on crab-eating macaque which is predicted to have undergone recent positive selection. We observed that the W98G substitution (which replaces a bulky tryptophan at the base of the pocket for the smallest residue, ancestral glycine) significantly increased binding affinity for endogenous lipids in crab-eating macaque, thus making it unlikely that swapping for DDM would occur (Figure 5A). This mutation appears to have similar effects in other genetic backgrounds as well, including humans (Figure 5A,B).

**Figure 5.**
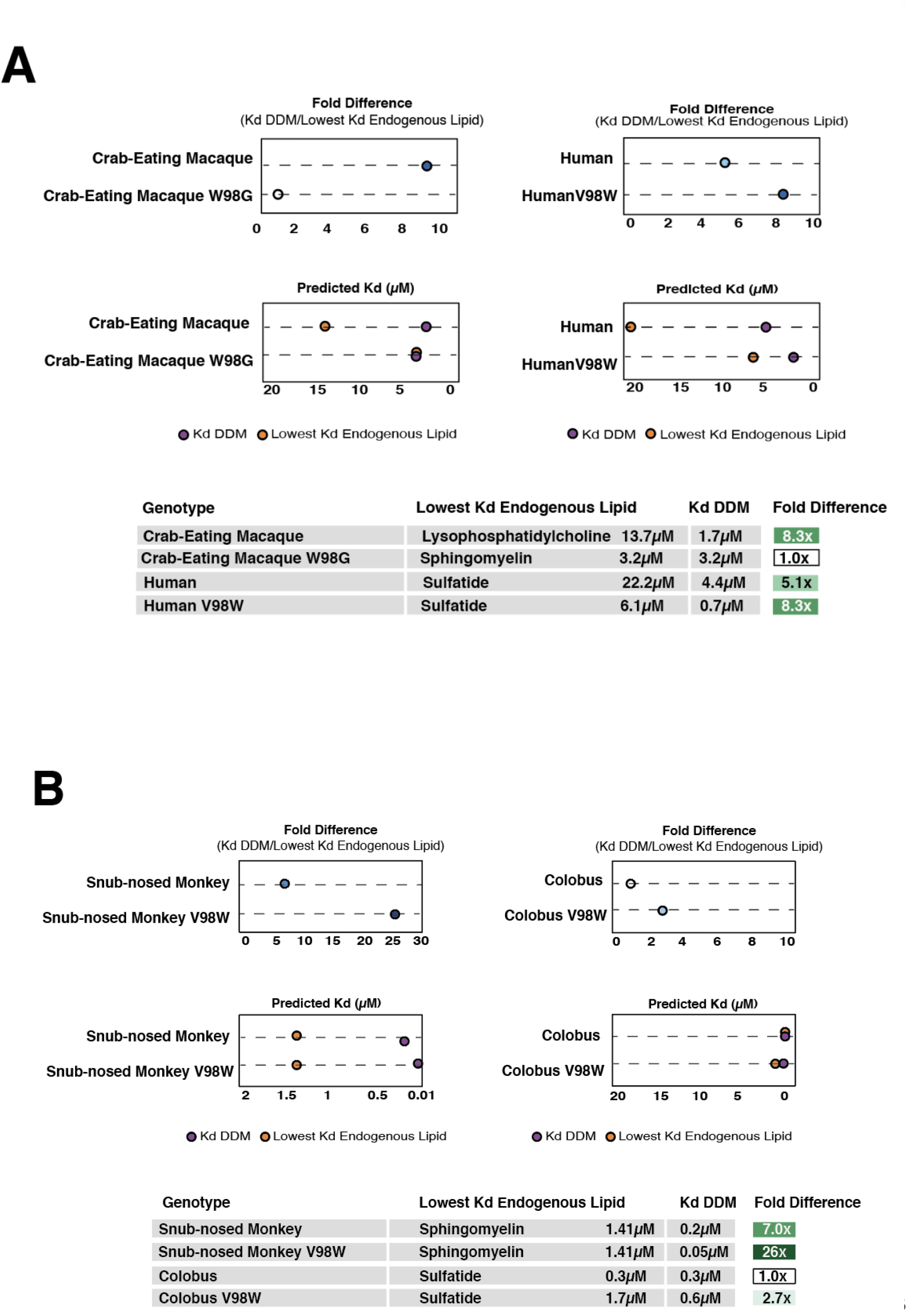
Rapidly-evolving positions in CD1a are sufficient to modulate predicted affinity for lipid antigens. **(A)** Mutation of site 98 to tryptophan in human CD1a (olive baboon and crab-eating macaque share this amino acid at this position) results in increased binding affinity to DDM, with overall fold increase between endogenous ligand and DDM. **(B)** Mutation of tryptophan at site 98 in crab-eating macaque to ancestral glycine results in higher binding affinity for all endogenous ligands tested, and loss of feasible lipid-swapping and DDM presentation. **(C)** Snub-nosed monkey, representing the consensus of the multiple sequence alignment at all sites under positive selection, also results in increased DDM binding when extant valine is mutated to tryptophan and a striking increase in spread between predicted endogenous and exogenous binding affinity. **(D)** Colobus, not predicted to swap endogenous ligand for DDM, also increases spread between binding affinities. In colobus, however, it is a decrease in affinity for endogenous lipid not increase in DDM affinity that is responsible for the fold change.

Analysis of the binding pose in crab macaque W98G bound to lysophosphatidylcholine shows the ligand buried in the pocket without exposed head group, a scenario which has demonstrated auto-immune effects in human CD1a (de Jong et al. 2014) (Figure 6A). This suggests an explanation for why the reduction in accessible pocket volume may be beneficial, both for lipid swapping mechanism and T-cell receptor ligand recognition.

**Figure 6.**
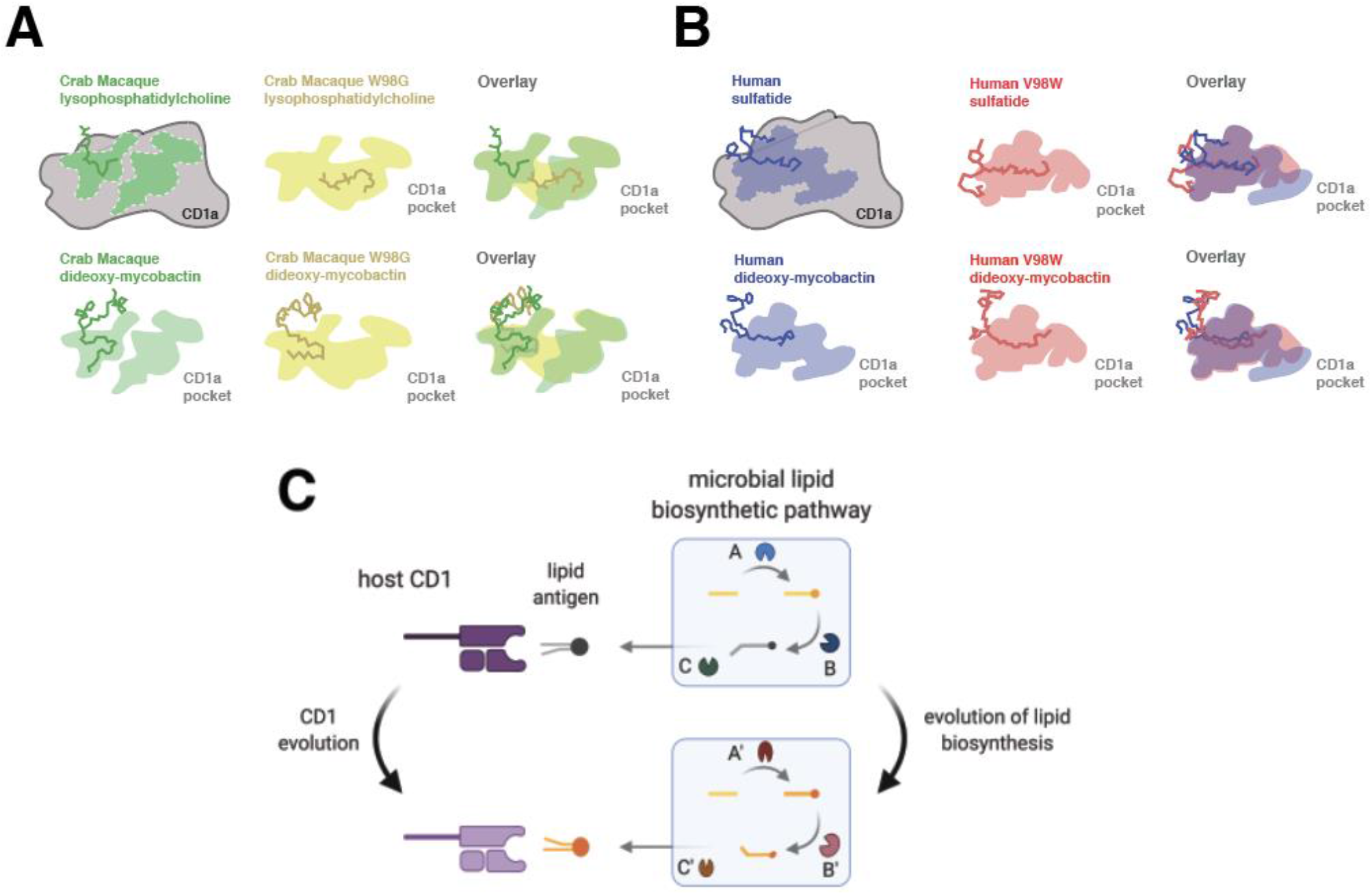
Conceptual framework for lipid-driven diversification of CD1 molecules. **(A)** Crab-eating macaque CD1a, which encodes a tryptophan in position 98, is predicted to lose the ability to present self-lipid lysophosphatidylcholine when this position is mutated to the consensus at this site, glycine. An overlay of the differences in pocket morphology shows how the tryptophan limits access to the deeper chambers of the pocket. (**B)** Humans possess a valine at position 98, which has been proposed to act as a molecular ruler limiting larger ligands access to the pocket. When this residue is mutated to a tryptophan *in silico*, further decreasing access to the deeper chambers of the pocket, the ability to swap out endogenous for exogenous ligand is improved, suggesting that a large hydrophobic residue in this position may be adaptive in the context of *M. tuberculosis* infection in primates. Cartoons were informed by analysis of Autodock Vina docking results analyzed in PyMol. **(C)** Proposed conceptual framework for lipid-driven evolution of CD1, resulting in accelerated evolution and rapid diversification of host immune receptors. Lipid biosynthesis pathways are complex and interdependent, thereby adding levels of complexity that may slow the rate at which pathogens can successfully alter new lipid antigens.

However, reduction in accessible pocket volume is not the only desirable feature for a functional CD1a molecule. Because the lipid-swapping mechanism requires higher binding affinity for exogenous lipids the pocket should be tuned to bind exogenous ligand with high affinity, which may require access to deeper pocket chambers. In the human V98W mutation, we noticed that the tail group accesses deeper regions of the pocket, which may partially explain the higher affinity for DDM seen in this model (Figure 6B).

To probe our system further, we used the genetic background of snub-nosed monkey to test broader-scale effects of mutations since it encodes primate consensus residues at positions with elevated dN/dS. We mutated seven sites that appear at the interaction interface to the ancestral sites at all loci, resulting in a protein that is not likely to swap endogenous ligand for DDM by our predictions (Supplemental Figure S6). Smaller effect mutations were found when introducing combinations of mutations in crab-eating macaque at position 114 where tyrosine appears to lower affinity for endogenous ligand and increases affinity for mycobacterial ligand slightly (Supplemental Figure S7). Taken together, several species are predicted to bind DDM with relatively high affinity but may not necessarily present exogenous antigen due to equally or greater affinity for endogenous lipid. This suggests that selective pressure may exist to decrease affinity for endogenous ligand in conjunction with increased affinity for exogenous antigens, resulting in increased effectiveness of CD1a-dependent immune responses. We observed that even single substitutions in rapidly-evolving sites substantially alter both endogenous and pathogen-derived lipid antigen recognition, providing further evidence for the functional impact of inter-species divergence in CD1a.

## Discussion

Overdominance has been proposed as an important driving force acting on MHC to promote diversity across the gene family (Hughes and Nei 1990). CD1 genes possess limited sequence variation within humans (Blumberg et al. 1995), which suggests overdominance is likely not a major factor shaping the evolution of this family. Rather, our observations of elevated dN/dS between CD1 orthologs and limited polymorphism within species are most consistent with a history of repeated selective sweeps driven by positive selection. Moreover, the patterns of divergence in CD1, with amino acid variation largely restricted to the MHC-like domain, supports the hypothesis that lipid antigen recognition and presentation are the functional drivers of this divergence. These patterns are also observed in MHC genes (which are also undergoing positive selection), with elevated ω at hotspots in the MHC antigen recognition site (Hughes and Nei 1990; Manlik et al. 2019). The electrostatic property variation in lipid ligands is found almost exclusively in the head-groups, with differences in the tail groups restricted to length and geometry of the hydrocarbon tails. As these tail groups have the most physical contact with CD1 binding pockets, it is logical that amino acids changes affecting the length and geometry of this pocket determine which hydrophobic chains can be accommodated. Patterns of evolution observed in CD1 could reflect a classical arms race in which host receptors and a subset of microbial antigens antagonistically co-evolve through time. Alternatively, selection in a fluctuating environment where the fitness benefit of recognizing a particular lipid antigen changes over time could also produce elevated patterns of divergence in CD1. Co-evolution between lipid antigens and host proteins would likely involve mutations in microbial genes responsible for lipid processing or modification (Figure 6C). Future studies could aid in determining how variation in lipid modifying genes shapes CD1-dependent immune responses to specific pathogens.

CD1 molecules possess the ability to bind and present hydrophobic antigens from a variety of pathogens, many of which likely remain to be described. It is notable, however, that the majority of CD1 antigens identified to date are derived from pathogenic mycobacteria including *Mycobacterium tuberculosis*, the causative agent of tuberculosis in humans. Tuberculosis remains a devastating human public health burden, recently accounting for more deaths due to infectious disease than any other single pathogen (Forrellad et al. 2013). It is tempting to speculate whether mycobacteria have indeed imposed particularly strong selective pressure on CD1 molecules during animal evolution. Given the limited effectiveness of the current tuberculosis vaccine (Gong, Liang, and Wu 2018; Schito et al. 2015), addition of CD1-targeted antigens in a next-generation vaccine could provide one avenue for increased efficacy (Gong, Liang, and Wu 2018). Functional characterization of diverse CD1 orthologs beyond humans may reveal whether detection of mycobacterial antigens is a widely conserved feature in this family, as well as possible routes to enhance CD1-mediated immunity against *M. tuberculosis*. Alternatively, evolution-guided development of synthetic lipid antigens that confer increased activation of CD1-responsive T-cells could provide an alternative strategy to enhance lipid-based vaccines.

Collectively our results suggest that natural selection has acted to decrease affinity for endogenous ligand while increasing affinity for exogenous antigen by CD1a. We observed that a single substitution can significantly alter the predicted effects of ligand binding affinity, with potential consequences for antigen presentation (Figure 5A,B). Notably, a major effect mutation identified in species undergoing episodic bouts of selection has the ability to reliably increase affinity for DDM and/or decrease the affinity of endogenous ligands by CD1a (Figure 5). Our analyses also indicate other residues determining binding pocket volume in human CD1a are undergoing repeated positive selection across primates. In particular, valine 28 has been reported to form a molecular barrier that acts as a size-limiting determinant for antigen binding (Zajonc et al. 2003). Notably, New World monkeys encode a smaller residue (glycine) at this position. Replacement of valine with glycine might be expected to expand the size of the binding pocket. However, our molecular modeling indicates that the binding pocket in the New World monkey lineages is predicted to be smaller than even the crab-eating macaque or mangabey, which bear an isoleucine and a threonine, respectively, at this same site. These observations suggest that molecular determinants of binding pocket volume and morphology are complex and influenced by a combination of variable amino acid substitutions. Additionally, the size of the binding pocket does not appear to correlate with feasibility of DDM presentation. These observations may reflect selection acting to produce a binding pocket that is able to swap out endogenous ligand without the need for a loading protein as seen in other CD1 paralogs. This feature enables CD1a to directly surveil the extracellular milieu for pathogen-associated molecules, a potential advantage compared to the other CD1 molecules which require lysosomal processing and accessory protein loading before antigen presentation can occur at the cell surface.

In order for a microbial pathogen to evolve alternative lipid antigen structures, mutations likely occur in genes responsible for synthesis or modification the lipid antigen. Mutations in processing and production of lipids will most likely have effects on steps of the biosynthesis pathways that are downstream of the mutated enzyme (Figure 6C). In the future it would be intriguing to test whether primate CD1a orthologs have evolved to detect other lipid types or variations of mycobactin derived from other pathogen sources. According to data from NIHTPR’s AceView (Thierry-Mieg and Thierry-Mieg 2006), gene expression of CD1a/c is exceptionally high in tissues in pig-tailed macaque, even though it is not predicted by our study to present mycobactin. Additionally, certain orthologs such as the marmoset CD1a exhibit very low gene expression (Thierry-Mieg and Thierry-Mieg 2006) and may be undergoing rapid birth-and-death evolution(Nei and Rooney 2005) and eventual pseudogenization. Such observations would be consistent with findings of dynamic CD1 gene duplication and loss across vertebrates (Nei and Rooney 2005). The significance of changes in endogenous lipid presentation will also be an area for important future investigation. Certain isoforms of sulfatide, for example, are associated with cancerous cells and when bound to CD1a can prime T-cells (Takahashi and Suzuki 2012). It has also been shown that presentation of endogenous ceramides by CD1d is associated with the ability to detect disease (Paget et al. 2019).

While we focused our molecular modeling and simulation studies on CD1a, comparable signatures of positive selection were identified in primate CD1c, CD1d and CD1e. Further investigation of these receptors and their cognate antigens would greatly advance our understanding of the importance for CD1 diversity in the evolution of vertebrate immunity. While lipids and other pathogen-derived macromolecules have long been appreciated as critical targets for host innate and adaptive immune responses, the potential for these factors to promote evolutionary conflicts with host species has been relatively unexplored. By combining comparative genetics and molecular modeling approaches, this study illuminates how lipid antigens have shaped fundamental features of primate immunity and the detection of globally devastating pathogens.

## Author Contributions

N.M.P. and M.F.B. conceived the study. N.M.P. performed all phylogenetic analyses, structural modeling, and ligand-binding simulations with assistance from H. A-Z. N.M.P. prepared the original manuscript and figures with assistance from M.F.B. All authors reviewed and edited the manuscript.

## Acknowledgements

We are grateful to Michael Harms, Edward Chuong, and members of the Barber lab, especially Emily Baker, for helpful discussions and feedback on the manuscript. This work was supported by National Institutes of Health grant R35GM133652 (to M.F.B.). N.M.P. was supported by a National Institutes of Health training grant T32HD07348. The funders played no role in study design, data collection, interpretation, or the decision to publish this study.

## Competing Interests

The authors declare no competing interests.

## Methods

### Phylogenetic analyses

A gene tree of primate CD1 was generated with PhyML (phylogenetics by maximum likelihood) with Bayes selection criterion and 1000 bootstraps (Z. Yang 2007). Between 18-21 primate cDNA sequences were aligned for each CD1a-e gene using MUSCLE (Supplemental Figure S8), sequences were trimmed manually as described in PAML manual, using the species phylogeny as reported by Perelman, 2007. Our CD1a dataset included all available nucleotide coding sequences (cDNA) for 19 primate species, with areas of ambiguity and stop codons removed. Positively selected sites for all CD1 genes were detected using the phylogenetic analysis by maximum likelihood (PAML) software package with F3X4 codon frequency model. Likelihood ratio tests compared pairs of site-specific models M1 with M2 (neutral and selection, respectively), M7 with M8 (neutral, beta distribution of dN/dS <1; selection, beta distribution dN/dS >1, respectively). Additional tests were performed which account for synonymous rate variation and recombination, including FuBAR (Murrell et al. 2013) and MEME (Murrell et al. 2012), using the HyPhy software package (Murrell et al. 2012; Murrell et al. 2013). We chose a stringent selection criteria for the sites we focused on in this study: PAML and FuBAR posterior probability of greater than or approximately equivalent to 0.9, MEME p-value of 0.1 or less. All sites analyzed (unless otherwise stated) fit these criteria under all three tests.

### CD1a Structural Predictions

The Eukaryotic Linear Motif resource (Kumar et al. 2020) (http://elm.eu.org/) was used to identify structural motifs from the primary amino acid sequence of CD1a. Primate CD1a structures were predicted with amino acid sequences submitted to I-TASSER server (J. Yang and Zhang 2015) (https://zhanglab.ccmb.med.umich.edu/) to generate structures for analysis using PyMol, primate structural alignment from 14 primate structures colored by conservation based on RMSD calculations from PyMol alignment (https://pymolwiki.org/index.php/Color_by_conservation) (Mura et al. 2010) CASTp for volume predictions, and for use in ligand docking simulations. To assess confidence in our structural predictions, isoform 1 of full-length human CD1a was analyzed (there are several crystal structures available for this molecule) with a C-score of −0.36. C-score values vary from −5, 2 with positive values indicating higher confidence, and only structures with C-values between −1 and 2 were used for analysis. Structures were analyzed using PyMol (The PyMol Molecular Graphics System, Version 2.0 Schrödinger, LLC). For binding pocket volume predictions, CASTp was used and the radius probe was set at 0.75 Å for each iTASSER-predicted structure submitted for analysis. The predicted volumes for all species were plotted using the Seaborn package in Python.

### Ligand Docking with AutoDock Vina

Re-docking with human CD1a was first performed to identify flexible residues required for all known ligands to re-dock in the same model. We calibrated our modeling by re-docking known ligands in our human CD1a iTASSER-predicted structure. According to our calculations, a comparison of CD1a crystal structures bound to the smallest and largest ligands (PDB ID 4X6D, 1XZ0) yields an RMSD of 1.23Å. This suggests there is flexibility in the CD1a pocket, supported by a number of hydrogen bonds between residues of the main CD1a binding domain alpha helices. Dorsal loop of alpha helix 2 was identified as required and made flexible in all primate CD1a structures analyzed. Receptors with amino acid side chains that occluded the binding pocket were also made flexible if not engaged in hydrogen bonds, and any of these additionally flexible residues (see Supplemental Data for details). AutoDockTools 1.5.6 (Trott and Olson 2010) was used to prepare the ligands and receptors for ligand docking. AutoDock Vina was run in the command line and docking results were analyzed in PyMol and plotted with the Python Seaborn package. A Python script was written to perform K_D_ calculations. Details of Vina settings including exhaustiveness, grid center and x,y,z coordinates are available in the Supplementary Data.

## Supplemental Figures

**Figure S1.**
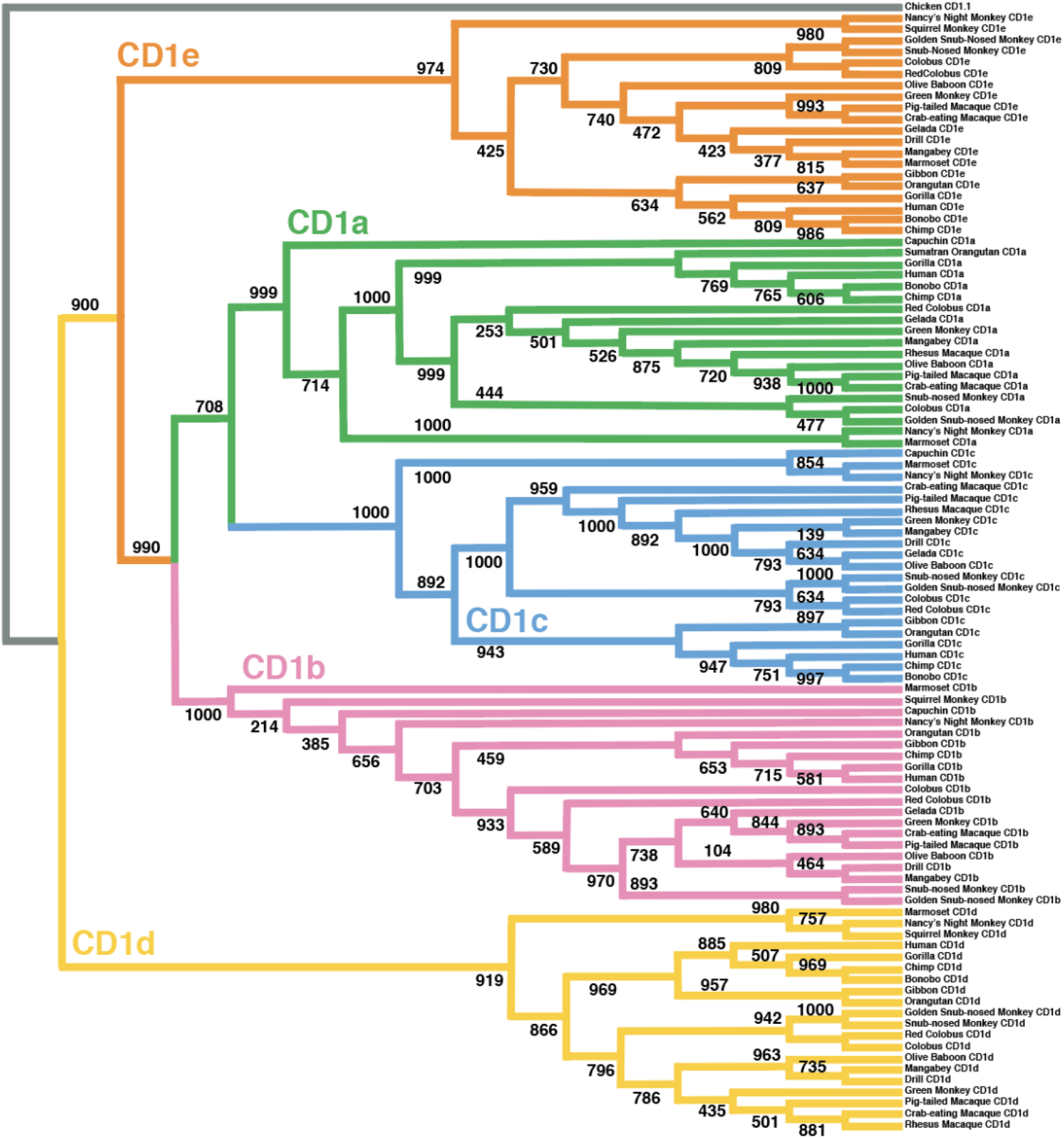
CD1 Primate Family Tree. Phylogenetic relationship of primate CD1 genes used in this study. Tree was generated in PhyML using maximum likelihood, 1000 bootstraps with ancestral Chicken CD1.1 as outgroup.

**Figure S2.**
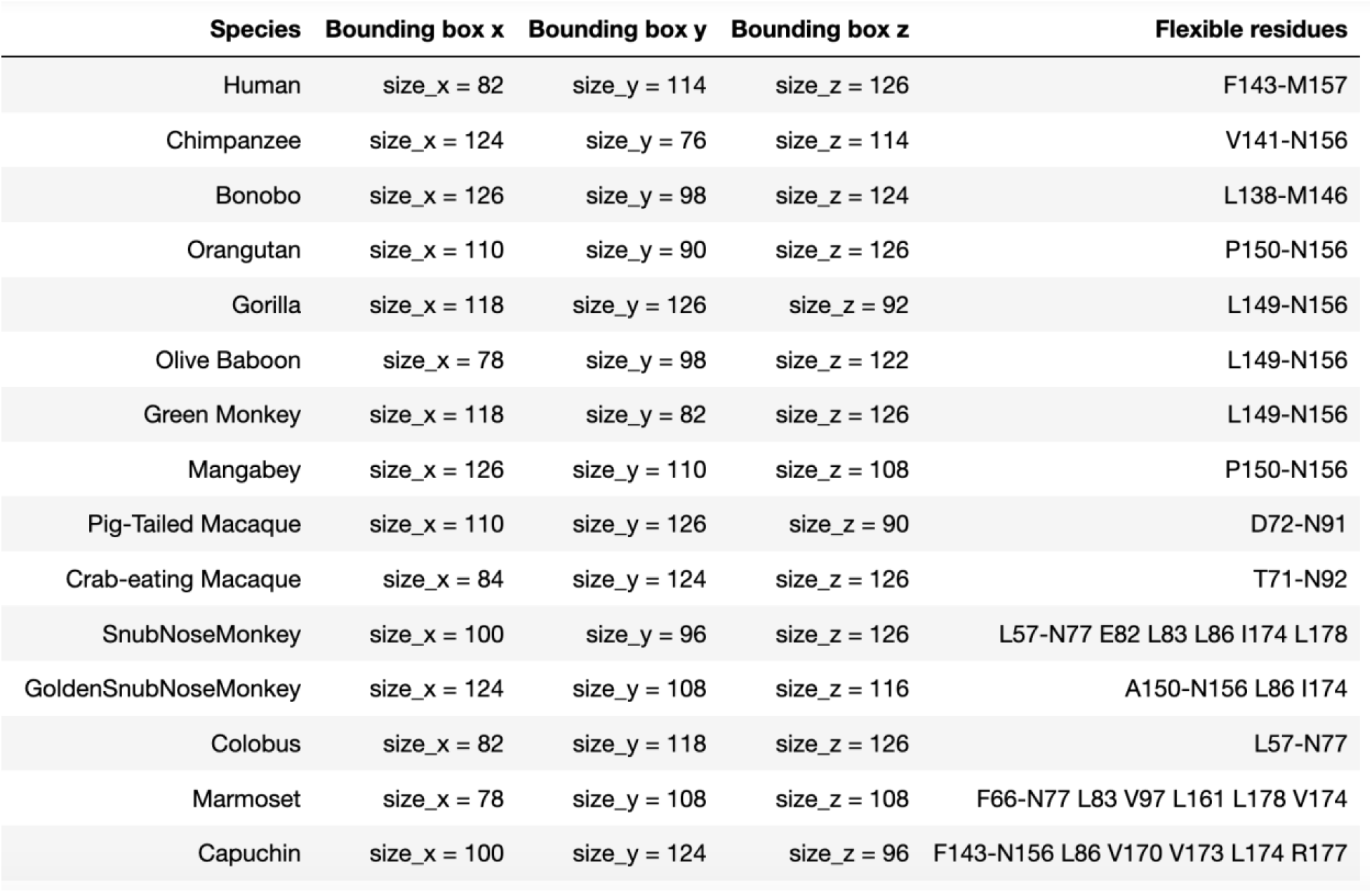
Vina Settings: Grid box parameters and flexible residues. Parameter settings for config file used to generate ligand docking models with Autodock vina, including residues set as flexible which correspond to the loop residues between alpha helices 1 and 2, and any occluding residues not engaged in hydrogen bonding at the portal entrance.

**Figure S3.**
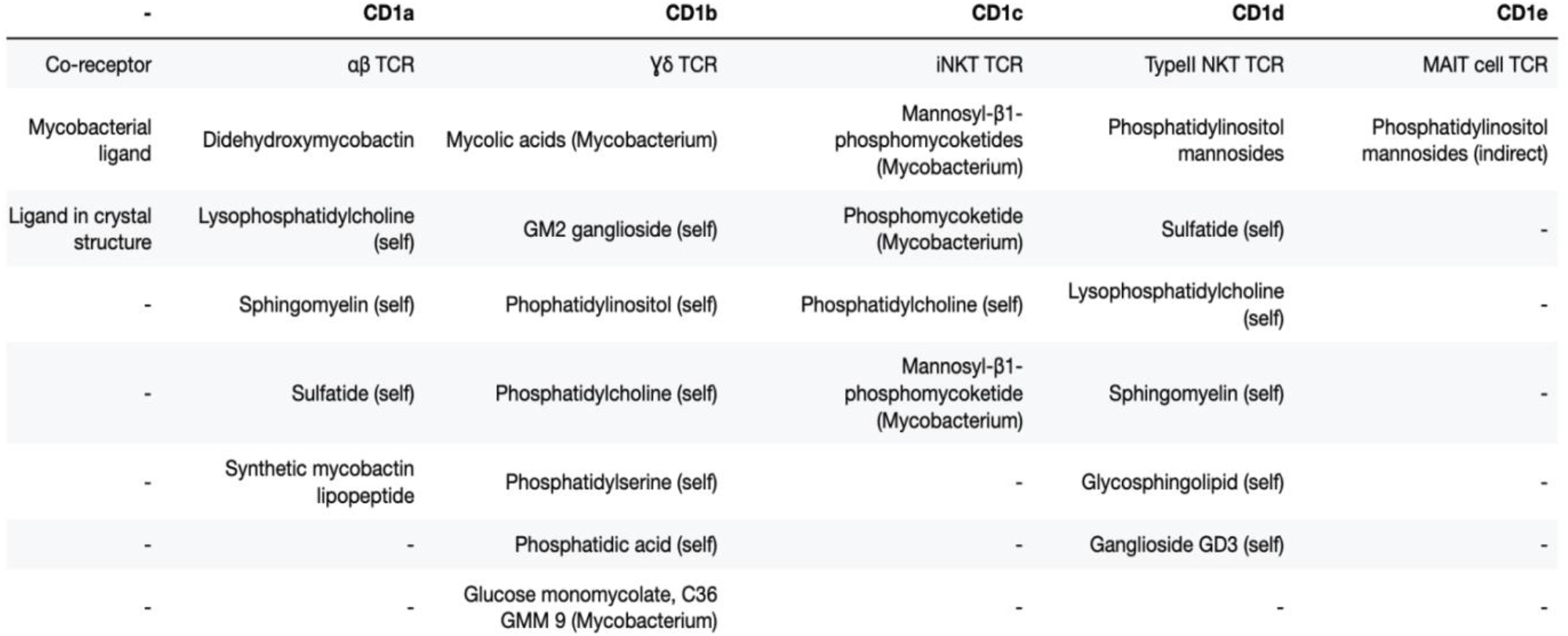
CD1 family members recognize a variety of mycobacteria-derived and endogenous lipids. Multiple mycobacterial lipids and lipoproteins are recognized by CD1 receptors, suggesting that this bacterial family known for exotic lipids has been interacting with the CD1 receptors across an extended timespan.

**Figure S4.**
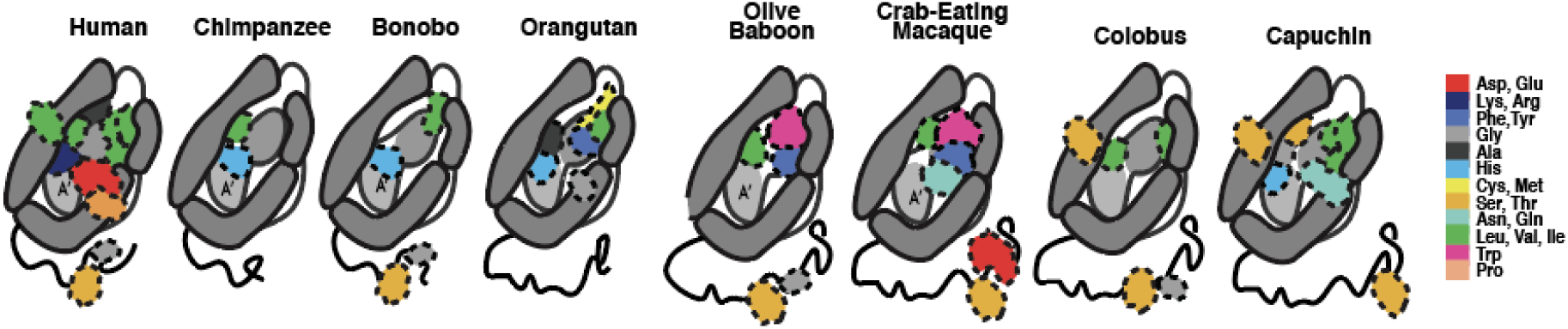
Sites under positive selection that differ from consensus, illustrated in a visual cartoon. Property differences in amino acids cluster around the portal and at the TCR interaction surface, while internal residues are mainly hydrophobic residues of varying sizes. Interestingly, the highly improbable substitution of the small glycine residue for the bulky hydrophobic tryptophan is common to Olive Baboon and Crab Macaque, both species predicted to have undergone multiple rounds of episodic adaptation in CD1a.

**Figure S5.**
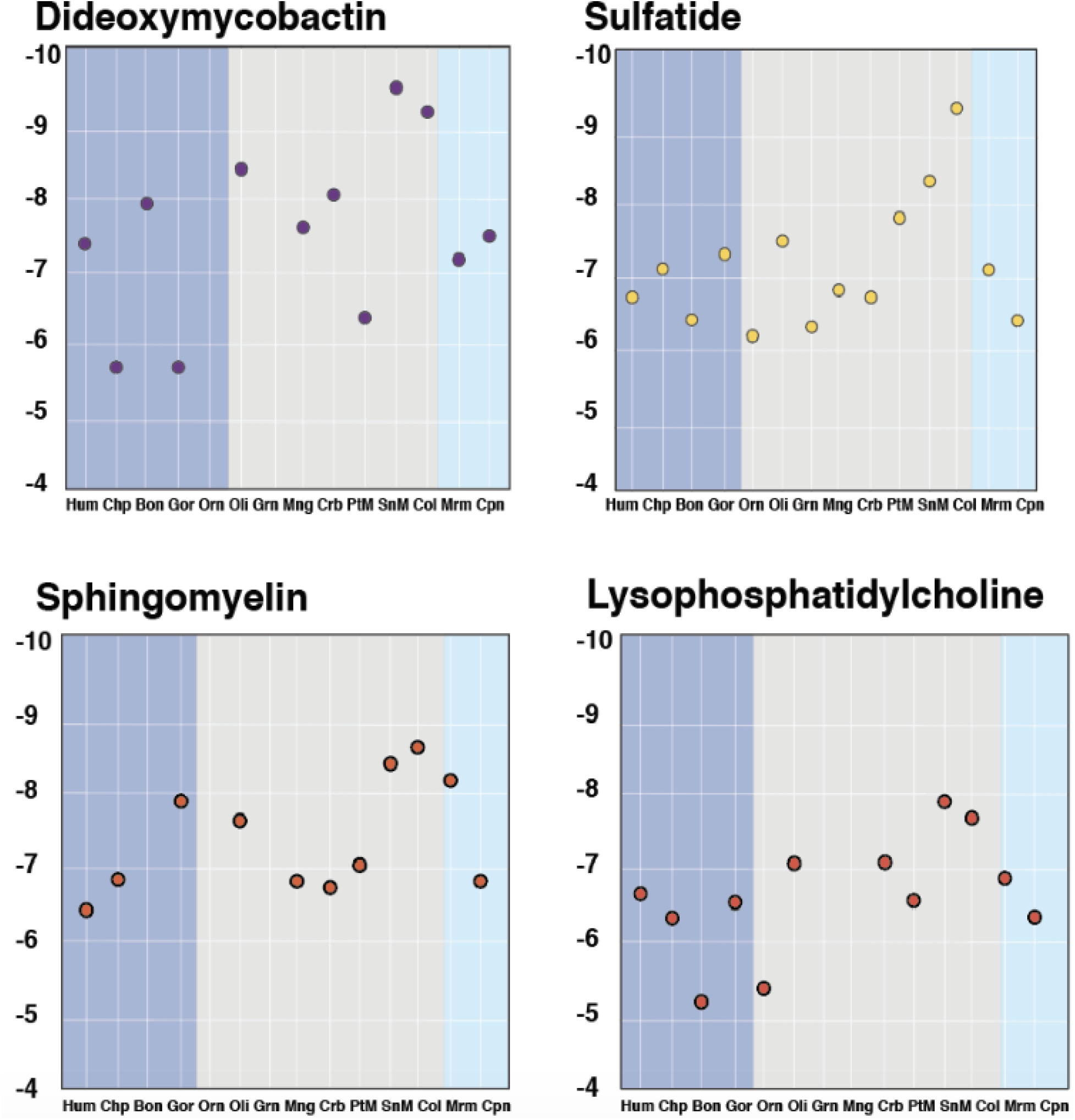
Plots of CD1a primate docking experiments by individual lipid. There appears to be little clustering when free energy values from ligand docking analyses are plotted by phylogenetic related-ness. **Dark blue**-Hominoids, **Grey**-Old World Monkeys, **Light blue**-New World Monkeys.

**Figure S6.**
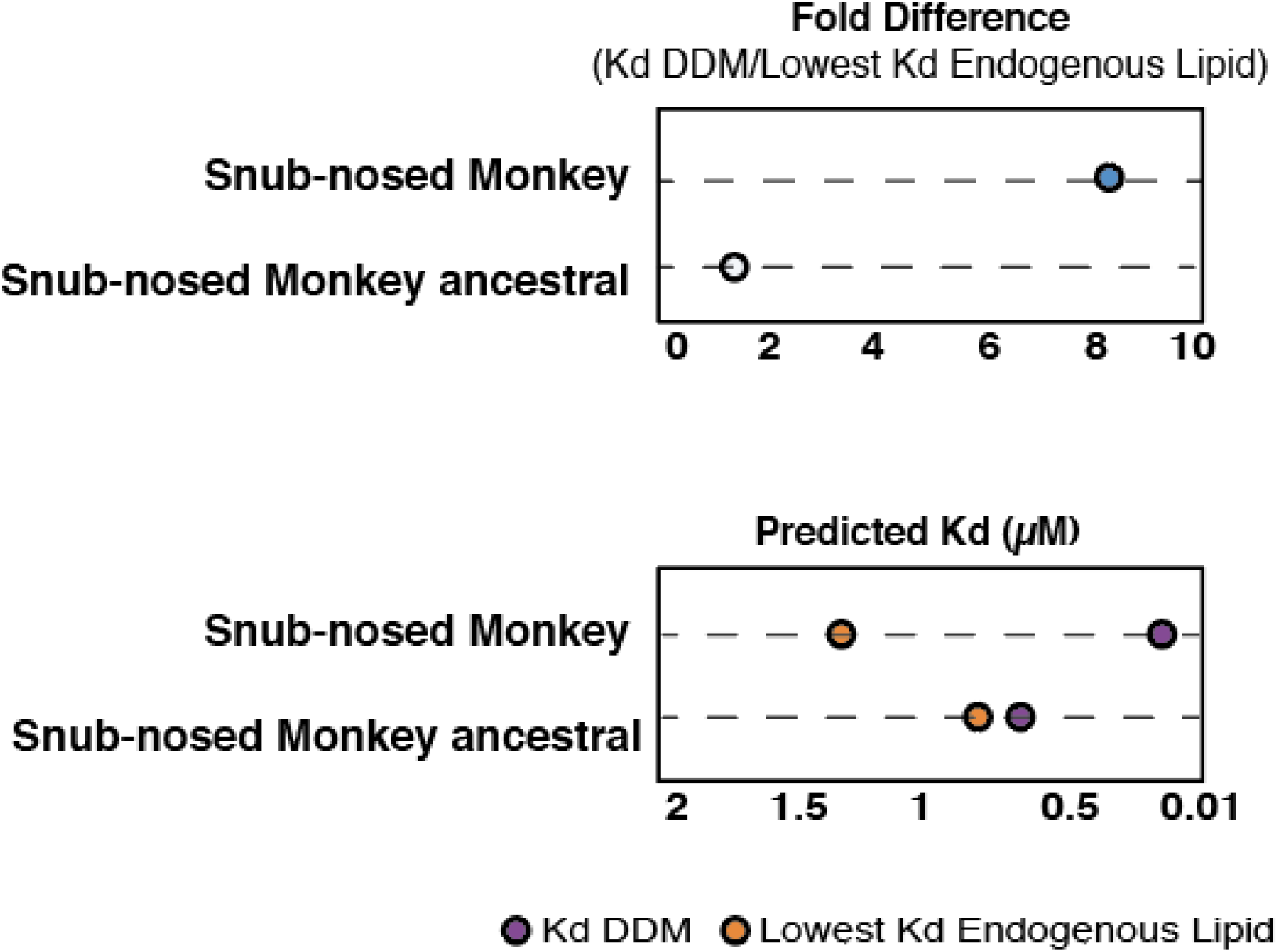
Snub-nosed monkey represents consensus at sites under selection. Ancestrally-predicted amino acids differ from consensus. Reversion of sites to ancestral state in Snub-nosed monkey background results in lower affinity for DDM and higher affinity for endogenous lipid.

**Figure S7.**
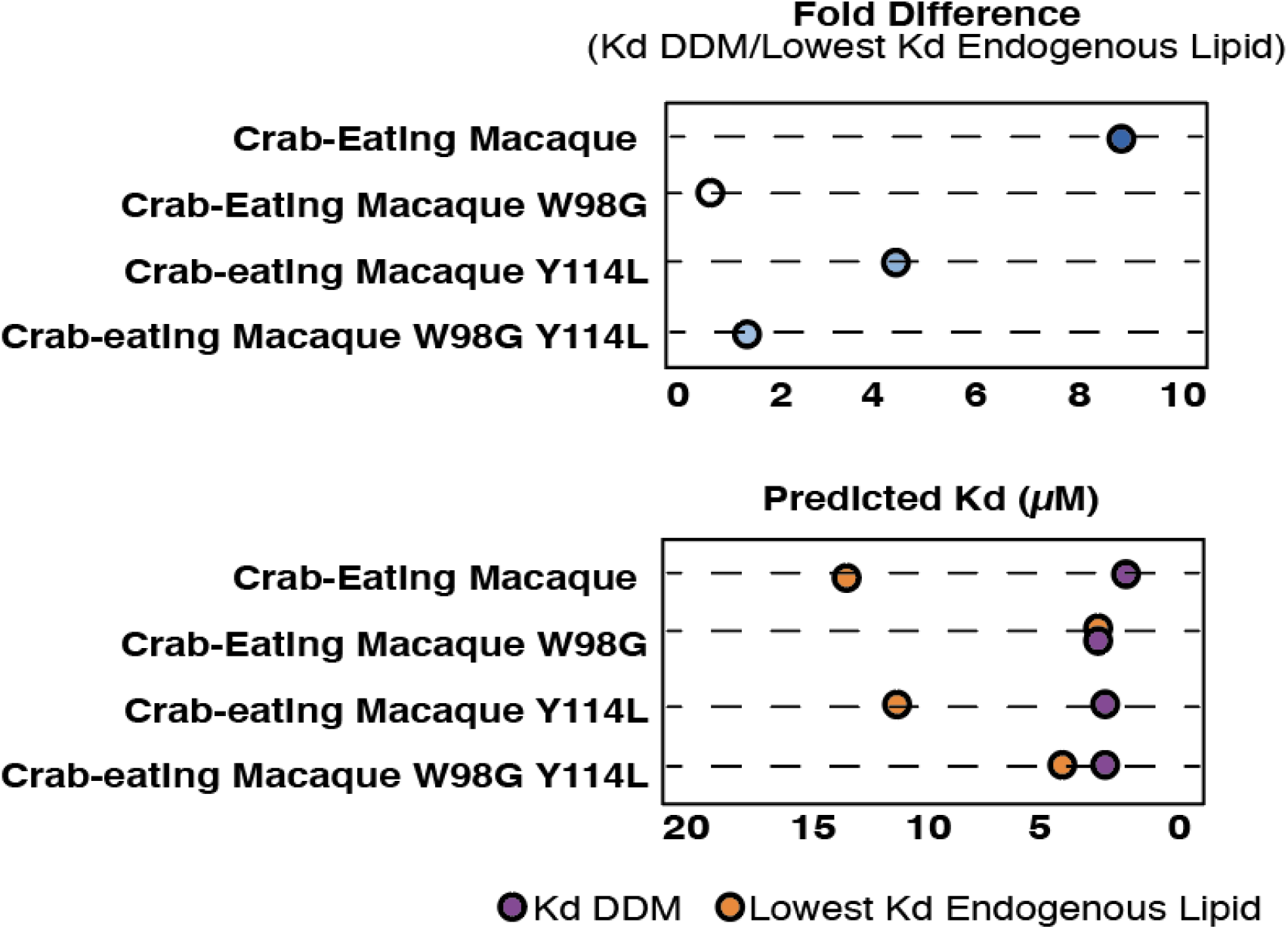
Additional amino acid changes confer minor increase in spread between endogenous and exogenous lipid. Small changes in affinity can be seen when other sites with high omega values are plotted in the Crab-eating macaque, such as site 114 which appears to have a similar effect to site 98, though smaller in magnitude, presumably due to the loss of bulky residue in the deeper chambers of the pocket.

**Figure S8. Species used for PAML analysis with NCBI accession identification numbers.**

**Primate CD1a, 18 species**

***Homo sapiens***, NCBI Reference Sequence: NM_001763.3; ***Macaca mulatta***, NCBI Reference Sequence: NM_001145818.1**; *Pan troglodytes***, NCBI Reference Sequence: XM_001169121.4; ***Macaca fascicularis***, NCBI Reference Sequence: XM_005595550.2; ***Papio anubis***, NCBI Reference Sequence: XM_003892864.5; ***Gorilla gorilla gorilla***, NCBI Reference Sequence: XM_004027022.3; ***Pan paniscus***, NCBI Reference Sequence: XM_024926560.2**; *Pongo abelii***, NCBI Reference Sequence: XM_002809984.3; ***Aotus nancymaae***, NCBI Reference Sequence: XM_012449747.1; ***Rhinopithecus bieti***, NCBI Reference Sequence: XM_017868910.1; ***Rhinopithecus roxellana***, NCBI Reference Sequence: XM_030935243.1; ***Colobus angolensis palliatus***, NCBI Reference Sequence: XM_011928485.1; ***Piliocolobus tephrosceles***, NCBI Reference Sequence: XM_023214151.2 ***Cebus capucinus* imitator**, NCBI Reference Sequence: XM_017503703.1; ***Callithrix jacchus***, NCBI Reference Sequence: XM_002764268.4; ***Chlorocebus sabaeus***, NCBI Reference Sequence: XM_007976629.1; ***Cercocebus atys***, NCBI Reference Sequence: XM_012093252.1; ***Macaca nemestrina***, NCBI Reference Sequence: XM_011769959.2

**Primate CD1b, 21 Species**

***Homo sapiens***, GenBank: AK303330.1**; *Papio anubis***, NCBI Reference Sequence: XM_003892865.5**; *Theropithecus gelada***, NCBI Reference Sequence: XM_025362870.1**; *Mandrillus leucophaeus*** NCBI Reference Sequence: XM_011978334.1**; *Cercocebus atys***, NCBI Reference Sequence: XM_012093255.1**; *Chlorocebus sabaeus***, NCBI Reference Sequence: XM_007976624.1**; *Macaca fascicularis***, NCBI Reference Sequence: XM_005541341.2**; *Macaca nemestrina***, NCBI Reference Sequence: XM_011769969.2 ***Rhinopithecus roxellana***, NCBI Reference Sequence: XM_010387232.2**; *Colobus angolensis palliatus***, NCBI Reference Sequence: XM_011957139.1**; *Piliocolobus tephrosceles***, NCBI Reference Sequence: XM_023214158.2**; *Nomascus leucogenys***, NCBI Reference Sequence: XM_003258651.4; ***Pan troglodytes***, NCBI Reference Sequence: XM_513909.5; ***Pongo abelii***, NCBI Reference Sequence: XM_002809978.3; ***Gorilla gorilla gorilla***, NCBI Reference Sequence: XM_004027024.3; ***Pan paniscus***, NCBI Reference Sequence: XM_003821008.2; ***Cebus capucinus imitator***, NCBI Reference Sequence: XM_017503705.1; ***Saimiri boliviensis boliviensis***, NCBI Reference Sequence: XM_003937886.2; ***Aotus nancymaae***, GenBank: AY605931.1; ***Callithrix jacchus***, NCBI Reference Sequence: XM_002760142.3

**Primate CD1c, 20 species**

***Homo sapiens***, Reference Sequence: NM_001765.3; ***Papio anubis***, NCBI Reference Sequence: XM_021925284.2; ***Theropithecus gelada***, NCBI Reference Sequence: XM_025393420.1; ***Mandrillus leucophaeus***, NCBI Reference Sequence: XM_011969349.1; ***Macaca mulatta***, NCBI Reference Sequence: NM_001145533.1; ***Macaca fascicularis***, NCBI Reference Sequence: XM_005595546.2, ***Macaca nemestrina***, NCBI Reference Sequence: XM_011769962.2, ***Rhinopithecus bieti***, NCBI Reference Sequence: XM_017868926.1**; *Rhinopithecus roxellana***, NCBI Reference Sequence: XM_010381246.2; ***Piliocolobus tephrosceles***, NCBI Reference Sequence: XM_023214153.2; ***Colobus angolensis palliates***, NCBI Reference Sequence: XM_011928487.1; ***Nomascus leucogenys***, NCBI Reference Sequence: XM_003258650.2; ***Pongo abelii***, NCBI Reference Sequence: XM_002809979.3; ***Pan paniscus***, NCBI Reference Sequence: XM_003821009.3; *Gorilla gorilla gorilla*, NCBI Reference Sequence: XM_019025012.2; ***Pan troglodytes***, NCBI Reference Sequence: XM_513908.6; ***Cebus capucinus imitator***, NCBI Reference Sequence: XM_017503701.1; ***Cercocebus atys***, NCBI Reference Sequence: XM_012093253.1; ***Callithrix jacchus***, NCBI Reference Sequence: XM_035279155.1

**Primate CD1d, 18 species**

***Homo sapiens***, NCBI Reference Sequence: NM_001766.4; ***Aotus nancymaae***, NCBI Reference Sequence: XM_012449750.2; ***Rhinopithecus roxellana***, NCBI Reference Sequence: XM_030935246.1; ***Cercocebus atys***, NCBI Reference Sequence: XM_012093246.1; ***Papio anubis***, NCBI Reference Sequence: XM_017948205.3; ***Macaca fascicularis***, NCBI Reference Sequence: XM_005541342.2; ***Macaca nemestrina***, NCBI Reference Sequence: XM_011769953.1; ***Chlorocebus sabaeus***, NCBI Reference Sequence: XM_007976632.1; ***Pongo abelii***, NCBI Reference Sequence: XM_024247050.1; ***Pan troglodytes***, NCBI Reference Sequence: NM_001071804.1; ***Pan paniscus***, NCBI Reference Sequence: XM_008974483.3 ***Macaca mulatta***, NCBI Reference Sequence: NM_001033114.2; ***Gorilla gorilla gorilla***, NCBI Reference Sequence: XM_019024988.2; ***Aotus nancymaae***, NCBI Reference Sequence: XM_012449750.2; ***Saimiri boliviensis boliviensis***, NCBI Reference Sequence: XM_010348509.1; ***Piliocolobus tephrosceles***, NCBI Reference Sequence: XM_023214147.2; ***Rhinopithecus bieti***, NCBI Reference Sequence: XM_017868929.1; ***Colobus angolensis palliates***, NCBI Reference Sequence: XM_011928483.1;

**Primate CD1e, 20 species**

***Homo sapiens***, NCBI Reference Sequence: NM_030893.4; ***Rhinopithecus roxellana***, NCBI Reference Sequence: XM_030935233.1; ***Macaca nemestrina***, NCBI Reference Sequence: XM_011769970.2; ***Macaca fascicularis***, NCBI Reference Sequence: XM_015455159.1*; **Cercocebus atys**,* NCBI Reference Sequence: XM_012093256.1; ***Chlorocebus sabaeus***, NCBI Reference Sequence: XM_007976621.1; ***Theropithecus gelada***, NCBI Reference Sequence: XM_025356228.1; ***Aotus nancymaae***, NCBI Reference Sequence: XM_012449741.1; ***Pongo abelii***, NCBI Reference Sequence: XM_003775532.3; ***Saimiri boliviensis boliviensis***, NCBI Reference Sequence: XM_003937882.2; ***Cercocebus atys***, NCBI Reference Sequence: XM_012093259.1; ***Gorilla gorilla gorilla***, NCBI Reference Sequence: XM_004027025.3; ***Pan troglodytes***, NCBI Reference Sequence: XM_513910.6; ***Pan paniscus***, NCBI Reference Sequence: XM_003821003.4; ***Nomascus leucogenys***, NCBI Reference Sequence: XM_003258652.4; ***Rhinopithecus bieti***, NCBI Reference Sequence: XM_017868912.1; ***Piliocolobus tephrosceles***, NCBI Reference Sequence: XM_023214159.1; ***Colobus angolensis palliatus***, NCBI Reference Sequence: XM_011957140.1; ***Papio anubis***, NCBI Reference Sequence: XM_021925303.2; ***Mandrillus leucophaeus***, NCBI Reference Sequence: XM_011978335.1;

## Notes

### Competing Interest Statement

The authors have declared no competing interest.

### Summary of Updates

The author list was entered in incorrect order.

